# Species differentiate water uptake depths within a narrow vertical zone in temperate grasslands

**DOI:** 10.64898/2026.11.09.765428

**Authors:** Sepideh Golshani, Tomas Hajek, Undine Schöllkopf, Johanna Harrison, Hassan Jafari, Barbora Rybova, Katja Tielbörger, Pierre Liancourt, Maria Majekova

## Abstract

Whether and under which conditions coexisting species in temperate grasslands differentiate their water uptake within the shallow soil layer they share is central to understanding hydrological niche differentiation in these ecosystems. In this field study, we quantified water uptake depth of 28 coexisting herbaceous species across 74 temperate grassland plots in three regions of Germany using oxygen stable isotopes and soil water content measured at multiple depths. Species consistently differed in water uptake depth across all regions, consistent with hydrological niche differentiation. This differentiation was largely confined to the upper 30 cm of soil and became stronger as overall soil moisture declined, with plants accessing water at greater depths in drier plots. In contrast, the vertical distribution of soil moisture was unrelated to either uptake depth or species differentiation. Despite pronounced species level variation, broad herbaceous functional classifications, including grasses versus forbs and fibrous versus taprooted species, showed substantial overlap in water uptake depth. Together, our results show that hydrological niche differentiation in temperate grasslands is expressed through fine scale differences among species within a shared shallow soil domain, becomes more pronounced under drier conditions, and is poorly captured by broad herbaceous functional classifications.

## Introduction

The precise mechanisms underlying how coexisting plant species differentiate in their use of resources, and how these differences contribute to the maintenance of biodiversity and ecosystem functioning, remain among the fundamental yet most debated questions in ecology. (MacArthur, 1955; Chesson, 2000; Silvertown, 2004). In particular, mechanisms related to belowground resource and space partitioning present a major challenge because belowground resource acquisition is difficult to observe and quantify directly. Thus, compared with aboveground interactions, empirical evidence for belowground niche differentiation under natural field conditions remains scarce (for review see Silvertown *et al*., 2015). Water, however, represents one resource along which coexisting plants may differentiate in their resource use, through differences in the depth or timing of water acquisition (Araya *et al*., 2011; Silvertown *et al*., 2015). Such differences can be interpreted as evidence of hydrological niche differentiation.

Temperate grasslands are a particularly informative system in which to test such differentiation, since approximately 80 to 90% of their roots occur within the upper 30 cm of soil (Kroon *et al*., 2012), and grasses predominantly acquire water from shallow soil layers (Feldman *et al*., 2023). This concentration of roots and water uptake within a shared soil volume can result in substantial overlap in water use among coexisting species, making consistent differentiation within this narrow vertical space particularly relevant. At the same time, water availability strongly constrains temperate grassland productivity (Prechsl *et al*., 2015), making changes in soil moisture particularly relevant under ongoing climate change (Lüscher *et al*., 2022; Kowalski *et al*., 2024). Whether and how species differentiate their water uptake depths as soils become drier is therefore increasingly important for understanding temperate grassland responses to changing water availability.

Water uptake depth represents a key functional dimension of belowground resource use, as it reflects where plants actually access soil water under field conditions. Importantly, water uptake depth is not equivalent to rooting depth; roots may be present substantially deeper than the soil layers contributing most strongly to water uptake, while uptake from a given root system can shift with soil water availability (Nippert *et al*., 2015; Guderle *et al*., 2018; Rickard *et al*., 2025; Kühnhammer *et al*., 2020). Water uptake therefore integrates several processes, including root distribution, root hydraulic properties, soil water availability and interactions with neighbouring plants. Consequently, although species can differ substantially in rooting architecture (Schenk and Jackson, 2002; Freschet *et al*., 2021), rooting architecture does not necessarily indicate the depth from which a plant is actively extracting water. This distinction is particularly important in herbaceous grasslands, where water uptake is often concentrated within the upper soil profile and several coexisting species may acquire water within the same relatively shallow soil volume (Bachmann *et al*., 2015; Prechsl *et al*., 2015; Tissink *et al*., 2025), despite substantial differences in their potential root architectures.

Previous studies in herbaceous grasslands provide mixed and strongly context dependent evidence for vertical differentiation in water use. Coexisting species can differ in their water uptake depths (O’Keefe *et al*., 2019; Nippert and Knapp, 2007), although uptake often remains concentrated within shallow soil layers and substantial overlap can occur among species and functional groups (Bachmann *et al*., 2015). Responses to declining water availability are similarly variable. Drought can shift water uptake towards deeper soil layers or induce species specific changes in uptake depth (Hoekstra *et al*., 2014; Tissink *et al*., 2025), whereas other grasslands show no general shift towards deeper uptake or even greater reliance on shallow water sources (Prechsl *et al*., 2015; Weides *et al*., 2024). Importantly, these responses are not necessarily shared among species. Under wetter conditions, coexisting species may rely on similar shallow water sources, whereas declining soil moisture may make differences in rooting distribution, hydraulic properties and plastic responses more consequential for where species acquire water (Nippert and Knapp, 2007). Lower soil moisture may therefore not only shift water uptake depth, but also increase differentiation among species.

Despite this context dependence, belowground water use in grasslands is frequently interpreted using broad herbaceous functional classifications, such as grasses versus forbs or fibrous versus taprooted species. Grasses are generally expected to rely on relatively shallow soil water through dense fibrous root systems, whereas forbs, particularly taprooted species, may have greater access to deeper soil layers (Schenk and Jackson, 2002; Hoekstra *et al*., 2014). However, rooting morphology describes the potential architecture for soil exploration rather than realized water acquisition. High variation among species within functional groups, together with environmentally driven plasticity in uptake depth, may therefore weaken correspondence between growth form, root system type and actual water uptake (Nippert *et al*., 2015; Guderle *et al*., 2018; Kühnhammer *et al*., 2020). Whether grasses and forbs, or fibrous and taprooted herbaceous species, consistently occupy distinct water uptake depths across natural grassland communities therefore remains unclear.

Stable oxygen isotopes have been widely used to infer plant water uptake depth by comparing isotopic signatures of soil water and plant xylem across vertical soil profiles (Ehleringer and Dawson, 1992; Ogle *et al*., 2004; Bachmann *et al*., 2015; Silvertown *et al*., 2015; Freyberg *et al*., 2020). Under dry intervals, evaporation enriches shallow soil water in ¹⁸O, generating a vertical gradient that can be traced in plant tissues with little additional fractionation during root uptake and transport (Barnard *et al*., 2006; Chen *et al*., 2020; Barnes and Turner, 1998). As a result, xylem δ¹⁸O provides a useful proxy for the soil depth from which plants acquire water, and is generally less sensitive to extraction artefacts than hydrogen isotopes (Ehleringer and Dawson, 1992; Orlowski *et al*., 2016; Gaj *et al*., 2017).

Here, we quantify species level water uptake depth of 28 species across 74 temperate grassland plots in three regions of Germany using oxygen stable isotopes to characterize water uptake of coexisting herbaceous species. We first test whether species differentiate their water uptake within the narrow vertical soil zone they share, providing evidence consistent with hydrological niche differentiation, and whether these differences are captured by broad herbaceous functional classifications based on growth form and rooting system. We then combine water uptake estimates with gravimetric soil water content measured at multiple soil depths to characterize overall soil moisture and its vertical distribution within each plot. We test whether lower overall soil moisture is associated with deeper water uptake and stronger differentiation among species. We further test whether species water uptake depth is related to the vertical distribution of soil moisture within the profile.

Specifically, we address three questions. First, do coexisting species differ in their water uptake depths across temperate grasslands, providing evidence consistent with belowground niche differentiation? Second, is lower overall soil moisture associated with deeper water uptake and greater species level differentiation, or are these patterns better explained by the vertical distribution of soil moisture? Third, do broad herbaceous functional classifications, specifically grasses versus forbs and fibrous versus taproot systems, explain the observed species differences in water uptake?

Understanding these patterns of species water uptake differentiation in temperate grasslands is therefore important for evaluating how belowground resource use may contribute to coexistence in these systems.

## Materials and Methods

### Study sites

The study was conducted on 74 permanent grassland plots across three regions (Table S1, Fig. S2), with 25 plots per region except for one missing plot in SCH. Plots spanned broad variation in bedrock, climate, and soil conditions and covered the full regional gradient of grassland management, including fertilization, mowing, and grazing (Fischer *et al*., 2010). The southern region, Schwäbische Alb (ALB), is situated in a limestone-dominated low mountain range in south-eastern Germany (460–860 m a.s.l.) within a UNESCO Biosphere Reserve. Soils in this region are mainly Leptosols and Cambisols, and mean annual precipitation ranges from 700 to 1000 mm. The central region, Hainich-Dün (HAI), lies in a hilly landscape within the National Park Hainich-Dün (258–550 m a.s.l.). Dominant soil types include Cambisols, Vertisols, and Stagnosols, and mean annual precipitation varies between 500 and 800 mm. The northern region, Schorfheide-Chorin (SCH), is in north-eastern Germany in the UNESCO Biosphere Reserve Schorfheide-Chorin and is characterized by gently undulating moraine terrain (3–140 m a.s.l.). This region encompasses a wide range of soil types, including drained Histosols, Luvisols, Gleysols, Cambisols, and Albeluvisols, and receives lower annual precipitation (480–580 mm) than the other two regions.

### Experimental design

Within each plot, a randomly located 1 × 1 m quadrat was selected for soil and plant sampling. To ensure sufficient aboveground biomass, particularly in grazed or mown plots, the quadrat and an approximately 2 m buffer zone were fenced one month before sampling.

### Soil and plant sampling

To estimate plant water uptake depth from natural soil δ¹⁸O gradients, a vertical isotopic stratification was allowed to develop during short dry periods (7–14 days), when evaporation enriches shallow soil layers in ¹⁸O (Schwinning, 2008; Barnes and Turner, 1998, Fig. S3). Because relying on natural dry spells was not feasible across regions, rainfall was excluded for 10–14 days using transparent roofs installed simultaneously on all plots in late May 2023. Roofs (2 × 2 m) were positioned above a central 1 × 1 m quadrat with a 0.5 m buffer zone and mounted on wooden frames (60–80 cm high) with open sides and a slight tilt to allow airflow and prevent rain intrusion. Microclimatic effects were assessed using TMS-4 data loggers placed under and 2 m outside the roofs in eight plots per region, recording soil moisture at 10 cm depth and air and soil temperature.

Sampling was conducted in early June 2023, 10–14 days after roof installation, and completed within one week across all regions. Soil cores (2 cm diameter) were taken to a maximum depth of 50 cm and sampled at 4, 8, 10, 12, 16, 32, and 48 cm, with finer resolution in the upper profile because most grassland water uptake occurs above 20 cm (Weides et al. 2024). Depth-specific soil samples (c. 2 g, root- and stone-free) were placed in 12 ml glass Exetainer® vials sealed with chlorobutyl septa, kept at 2–4 °C in the field, and stored at −20 °C. To minimize moisture loss, the auger was wrapped in cling film during coring and opened only at target depths.

Plant material was collected within 50 cm of each soil core from locally abundant species (5–12 species per plot, Table S2, Fig. S2). Root crowns were sampled because they best represent source water in herbaceous plants (Barnard *et al*., 2006). For each species, 0.15–0.5 g fresh mass was collected, pooling multiple individuals when necessary. Samples were cleaned of soil and green tissue, placed in 12 ml Exetainer® vials, and stored as described for soil.

### Water extraction and oxygen stable isotope analysis

Water was extracted from soil and plant samples by cryogenic vacuum distillation. Paired 12 ml Exetainer® vials were connected via threaded adapters and evacuated to c. 100 Pa, then heated to 100 °C while vapor was condensed in a liquid-nitrogen trap. Extraction lasted 1.5–2.5 h per batch (up to 48 samples). Extraction efficiency was assessed by repeated weighing before and after distillation and after reheating to detect leakage or incomplete recovery. Samples showing anomalous isotope values or mass loss were excluded. Up to 250 µl of extracted water was transferred to airtight autosampler vials for isotope analysis.

Oxygen stable isotope ratios (δ¹⁸O) of extracted water were measured using an isotope-ratio mass spectrometer (DELTAplus XL) coupled to a high-temperature conversion elemental analyzer (TC/EA; ThermoFinnigan, Bremen, Germany). A laboratory standard was analysed after every ten samples to monitor analytical consistency. Isotope values are reported in per mil (‰) relative to Vienna Standard Mean Ocean Water (VSMOW). Oxygen isotopes were used instead of hydrogen isotopes because they are less sensitive to extraction artefacts and provide more reliable estimates in herbaceous tissues (Chen et al. 2020; Weides et al. 2024).

### Soil moisture content

Gravimetric soil water content, hereafter soil moisture, was calculated for each depth specific soil sample as:

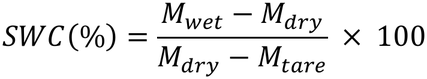

where (*M_wet_*) is the mass of the vial plus wet soil, (*M_dry_*) is the mass of the vial plus dry soil, and (*M_tare_*) is the empty vial mass. Overall soil moisture, representing plot level soil wetness, was defined as the median gravimetric soil water content across sampled depths within each plot. Vertical soil moisture distribution was calculated as the Pearson correlation between sampling depth and gravimetric soil water content within each plot. Positive values therefore indicate increasing soil moisture with depth, whereas negative values indicate higher soil moisture towards the soil surface.

### Statistical analyses

δ¹⁸O values were modelled as a function of log-transformed soil depth using mixed-effects models for each region (ALB, HAI, SCH) (Weides *et al*. 2024), we included the plot-specific random intercepts and slopes to account for spatial heterogeneity among soil profiles. The fitted models were then used to estimate plant water uptake depth by inserting the δ¹⁸O value measured in the root crown into the modelled soil isotope profile from the plots (Fig. S1).

Hydrological niche differentiation was assessed using Kruskal–Wallis tests (kruskal.test, R base), reported as χ² statistics. Species-level variance in uptake depth was quantified using intraclass correlation coefficients (ICC) derived from intercept-only mixed-effects models with species and plot as random effects (lmer, package *lme4*).

To test whether soil moisture was associated with water uptake depth, we fitted a linear mixed effects model with log transformed predicted uptake depth as the response, overall soil moisture, vertical soil moisture distribution and region as fixed effects, and species and plot as random intercepts.

For water uptake differentiation among coexisting species within plots, mean pairwise differences in log water uptake depth were analysed at the plot level using a linear model with overall soil moisture, vertical soil moisture distribution and region as predictors. As a sensitivity analysis, we repeated both soil moisture models with land use intensity (LUI) included as an additional predictor to assess whether the observed moisture relationships were independent of grassland management. Full model results and LUI calculation details are reported in the Supporting Information (Table S4)

Overlap in water uptake depth between growth forms (grasses vs. forbs) and root system types (fibrous vs. taproot) was quantified for each region using kernel density estimation. Overlap was calculated as the area of intersection between density functions (package overlapping; Pastore, 2018), following established approaches (Mason *et al*., 2011; Le Provost *et al*., 2021). In addition, plot-level overlap was assessed for subsets with sufficient representation, restricting analyses to plots containing at least three species per group.

All analyses were conducted in R version 4.3.1 (R Core Team 2023).

## Results

### Species-level differentiation in water uptake depth

Plant water uptake depths differed significantly among coexisting species within each of the three regions (Fig. 1). Species-level differences were detected in ALB (χ² = 48.69, p = 0.008), HAI (χ² = 62.98, p = 0.0011), and SCH (χ² = 36.83, p = 0.0004). Most species accessed water in the upper soil layers up to the depth of 10 cm in ALB, 20 cm in HAI, and 30 cm in SCH.

**Figure 1.**
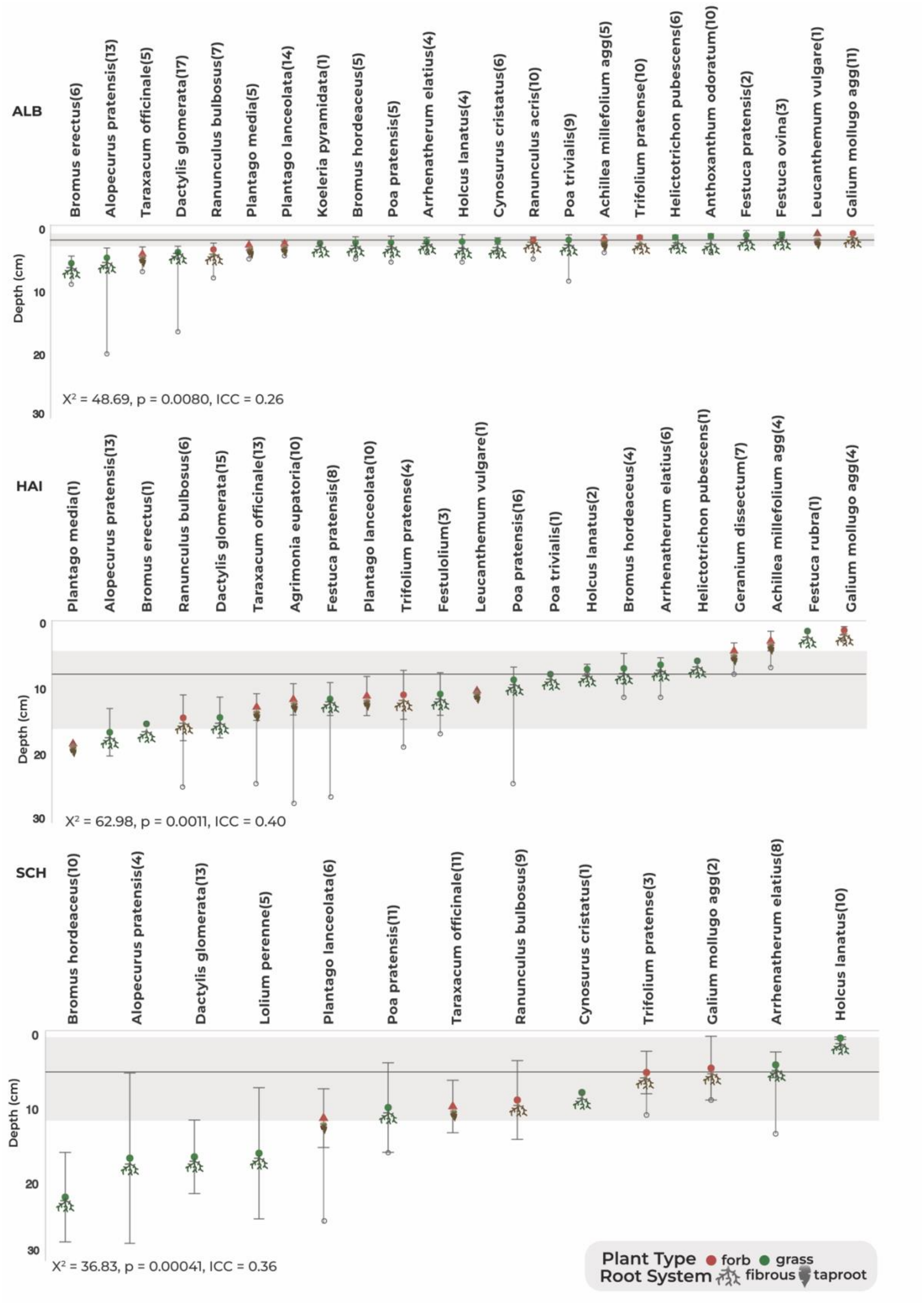
Belowground niche partitioning of plant species across the three regions (ALB, HAI, SCH). Predicted water uptake depth (mean ± SE) per species based on δ¹⁸O profiles. Points represent species means, horizontal bars indicate ± SE, and open circles show maximum observed uptake depth. Shaded bands show the interquartile range (25–75%) of uptake depths across all individuals within each region, with the horizontal line indicating the median (“focused uptake” zone). N in brackets indicates the number of plots in which each species was sampled. Symbols reflect plant type (color) and root system (shape). χ² and p-values indicate overall differences among species in predicted uptake depth within each region (Kruskal–Wallis test). Intraclass correlation coefficients (ICC) indicate the proportion of variance in uptake depth explained by species and plot within each region.

Species identity accounted for a substantial proportion of the total variation in uptake depth, explaining 26% of the variance in ALB, 40% in HAI, and 36% in SCH (intraclass correlation coefficients). Species differed in their maximum predicted uptake depths and in their positioning relative to the region-specific depth zone where water uptake was most concentrated (Fig. 1). Species-specific regional differences are presented in Table S3 and Fig. S4.

### Soil moisture and species-level differentiation in water uptake depth

Overall soil moisture was strongly associated with water uptake depth across grasslands (Fig. 2a). Across individual observations, water uptake depth increased as the overall soil moisture of the corresponding plot declined (β = −0.033, p < 0.001), whereas the vertical distribution of soil moisture within the profile was not significantly associated with uptake depth (β = 0.116, p = 0.521; Fig. 2c).

**Figure 2.**
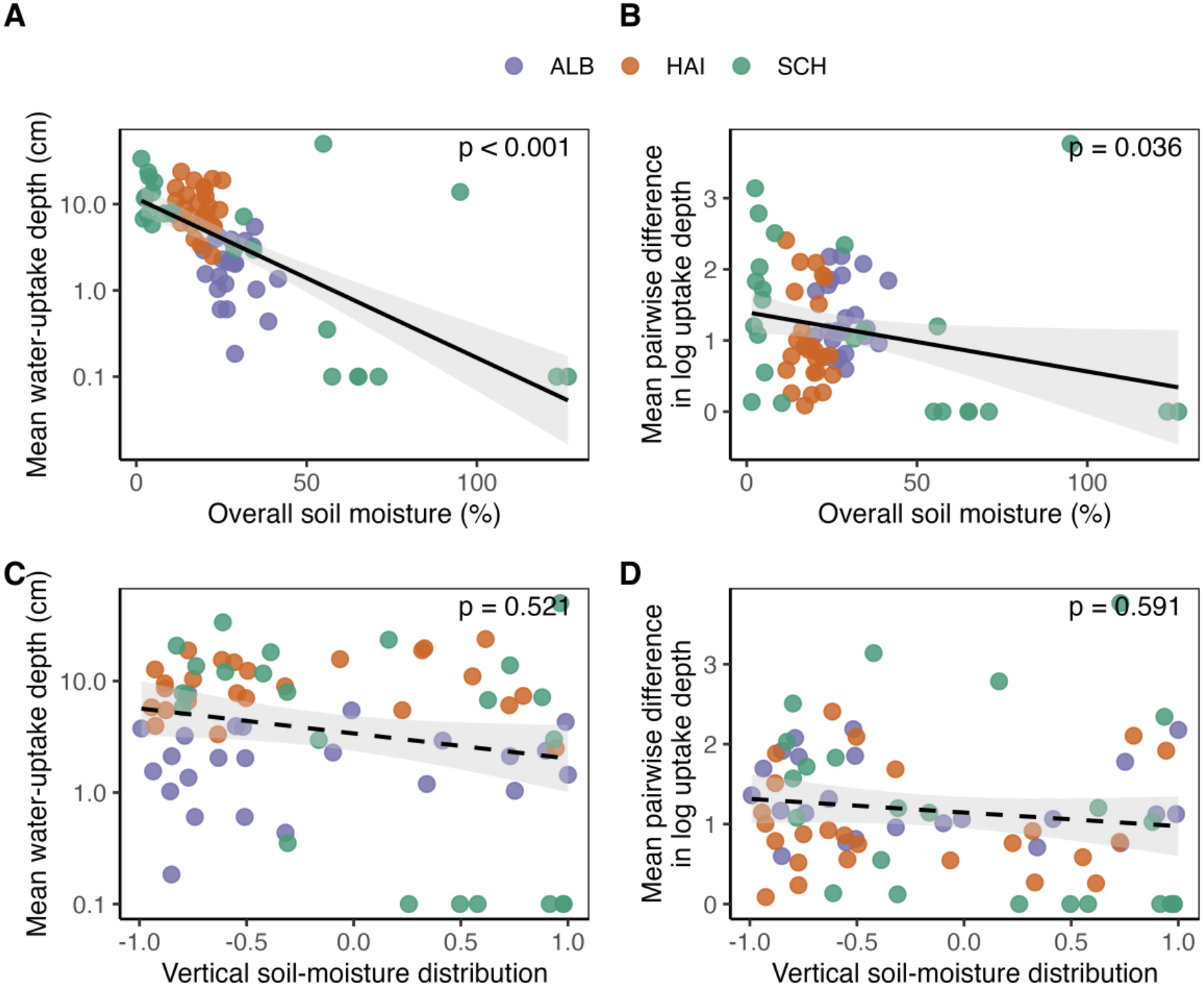
Effects of soil moisture on water uptake depth and species differentiation. Relationships between overall soil moisture and (A) mean water uptake depth and (B) mean pairwise differences in log water uptake depth among coexisting species, and between vertical soil moisture distribution and (C) mean water uptake depth and (D) mean pairwise differences in log water uptake depth. Points represent plots and colors indicate the three regions (ALB, HAI, SCH). Black lines show fitted relationships with 95% confidence intervals. P values are from models including overall soil moisture, vertical soil moisture distribution and region simultaneously.

Differentiating water uptake depth among coexisting species in a plot were greater at lower overall soil moisture (β = −0.009, p = 0.036; Fig. 2b), whereas they were not significantly related to the vertical distribution of soil moisture (β = −0.081, p = 0.591; Fig. 2d).

These relationships remained when land use intensity was included in the models, while LUI itself was not significantly associated with water uptake depth or mean pairwise differences among species (Table S4).

### Overlap in water uptake depths between growth forms

Despite these species-level differences, substantial variation within growth-form categories (forbs *vs* grasses) resulted in high overlap in their water uptake depth distributions within each region (Fig. 3a; 0 = no overlap, 1 = complete overlap). Overlap values were 0.89 in ALB, 0.96 in HAI, and 0.84 in SCH. In ALB, uptake distributions of both growth forms were strongly skewed towards the shallowest soil layers, whereas in HAI and SCH distributions were broader and less skewed. In SCH, grasses showed a slight tendency towards shallower uptake than forbs, but overlap remained high.

**Figure 3.**
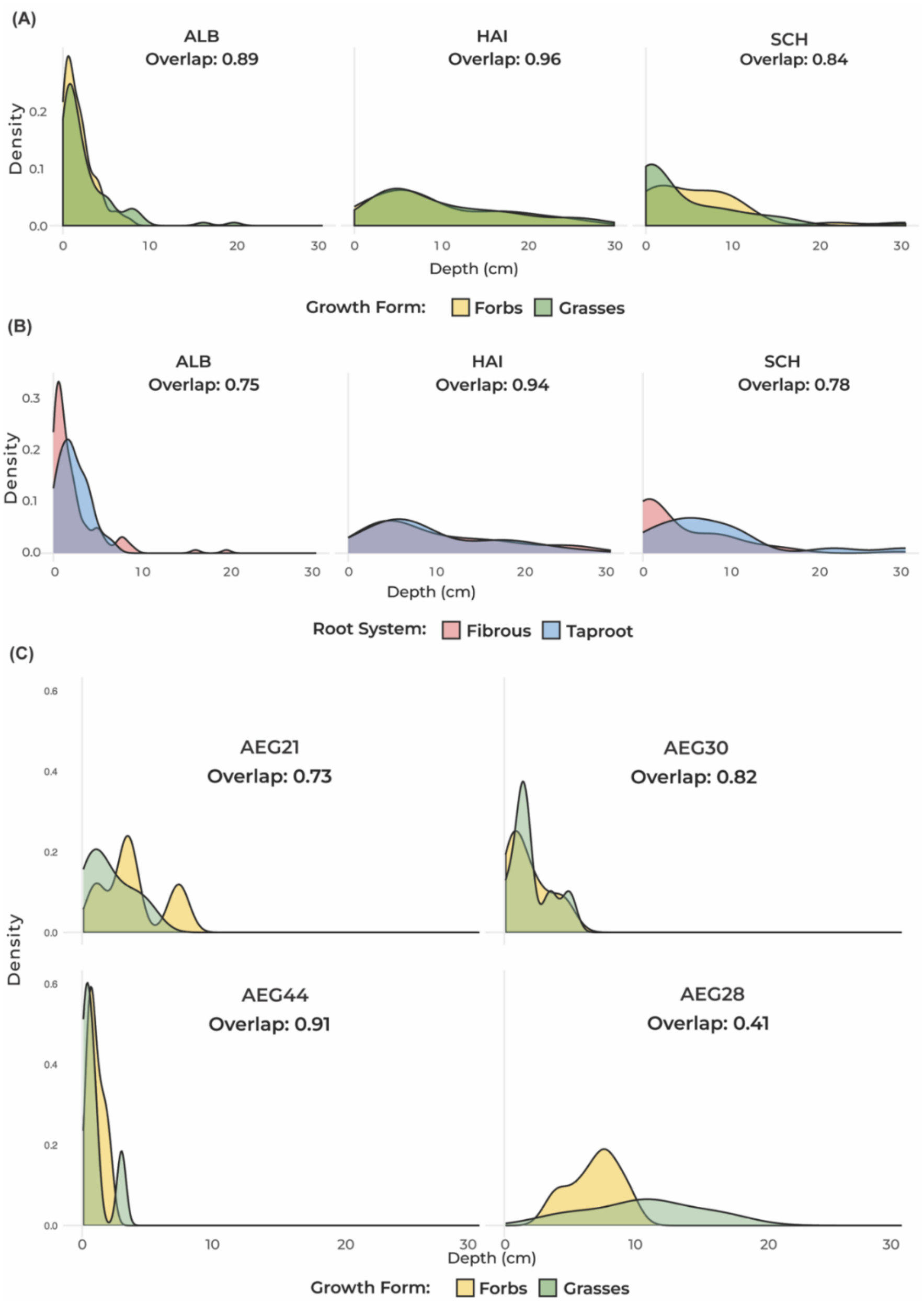
Overlap between growth forms and rooting systems. Overlap in water uptake depth between growth forms (grasses and forbs) (A) in the three regions and (C) in selected plots, and (B) between rooting systems (fibrous and taproot) in the three regions. Higher values indicate higher overlap with a maximum overlap being 1.0.

At the plot scale, only communities containing comparable numbers of grasses and forbs were considered. Despite greater heterogeneity in species-level uptake patterns, overlap between grasses and forbs remained high in most plots (Fig. 3c), ranging from 0.73 to 0.91.

### Overlap in water uptake depths between rooting system types

Similarly, substantial variation among species within rooting system types (fibrous *vs* taproot) led to consistently high overlap in their water uptake depth distributions across regions (Fig. 3b). Overlap values were 0.75 in ALB and 0.78 in SCH, reflecting a modest tendency for taprooted species to access slightly deeper soil layers in these regions, whereas overlap was nearly complete in HAI (0.94).

## Discussion

### Evidence for hydrological niche differentiation within a narrow vertical zone

Our results show that coexisting species consistently differed in their water uptake depths despite substantial overlap within a shared shallow soil zone. Across all three regions, this species differentiation was significant but largely confined to the upper 30 cm of soil.

This pattern is particularly relevant in temperate grasslands, where roots and water uptake are concentrated in the upper soil profile and substantial overlap in belowground resource use can therefore be expected. Previous studies have shown that coexisting grassland species can differ in their water sources or uptake depths (Nippert & Knapp, 2007; Hoekstra et al., 2014; O’Keefe et al., 2019), while studies comparing water use across shallow and deeper soil layers have also reported predominantly shallow uptake and substantial overlap (Bachmann et al., 2015). Given this strong concentration of water use near the surface, our sampling design used finer resolution in the upper soil profile, allowing us to resolve species differences within this narrow shared zone.

Several lines of evidence indicate that these species differences can be interpreted as hydrological niche differentiation. First, species differed significantly in uptake depth across all three regions, and species identity explained a substantial proportion of variation in uptake depth, indicating that these differences were not attributable solely to variation among plots. Second, these differences occurred within the same narrow soil domain rather than through broad separation between shallow and deep water sources. Third, mean pairwise differences in water uptake depth increased as overall soil moisture declined, indicating stronger species differentiation under drier conditions. Together, these patterns provide field evidence that water uptake depth represents a hydrological niche axis in temperate grasslands. However, establishing whether this hydrological niche differentiation contributes to stable coexistence would require direct tests linking differences in resource use to species interactions and population dynamics (Chesson, 2000; Levine et al., 2008; HilleRisLambers et al., 2012).

### Drier soil strengthens species differentiation in water uptake

Plants not only accessed water at greater depths as the overall soil moisture of their corresponding plots declined, but differences in water uptake depth among species also increased in drier plots. Water uptake can respond dynamically to soil water conditions and shift independently of root distribution (Guderle et al., 2018; Kühnhammer et al., 2020). In experimental drought studies have also reported shifts in grassland water uptake towards deeper soil layers without a corresponding increase in deep rooting (Tissink et al., 2025). However, drought responses in water uptake depth are variable; other temperate grasslands showed no general shift towards deeper uptake or even greater reliance on shallow water sources (Prechsl et al., 2015; Weides et al., 2024). Our results therefore extend this variable experimental evidence across a broad set of field communities, where lower overall soil moisture was consistently associated with deeper water uptake.

Importantly, species differentiation increased as plots became drier. Similar patterns have been reported at smaller experimental scales, where differences in water acquisition among species became more pronounced as shallow water availability declined (Nippert & Knapp, 2007; Hoekstra et al., 2014). This stronger plot level differentiation under lower water availability may indicate greater expression of hydrological niche differentiation as water becomes limiting.

Our results however do not support a simple explanation in which water uptake depths merely follows the vertical distribution of wetter soil layers. The vertical distribution of gravimetric soil moisture was unrelated to both uptake depth and differences among species water uptake depths within a plot, whereas overall soil moisture was associated with both. Similar mismatches among soil moisture, root distribution and uptake have been observed in herbaceous plants and grassland communities (Guderle et al., 2018; Kühnhammer et al., 2020). Thus, variation in overall soil moisture was more closely associated with water uptake depths than the measured vertical distribution of soil moisture.

Gravimetric soil water content reflects soil water amount rather than its availability to plants, which also depends on soil water potential and soil properties (Jones, 2007). Our measurements represent conditions at sampling and do not capture short term moisture dynamics (Beyer et al., 2020). In addition, the vertical moisture index summarizes the overall trend with depth and may miss more complex variation among soil layers. Within these limitations, vertical soil moisture was not associated with uptake depth or species differentiation.

### Broad herbaceous functional classifications poorly predict water uptake depths

Despite consistent species level differentiation, growth form classifications explained little of the observed variation in water uptake depth. Grasses and forbs exhibited high overlap in uptake distributions in all regions, even where their depth profiles differed slightly. Thus, the species differences detected in realized water uptake did not scale directly to a broad separation between grasses and forbs. This finding challenges the common expectation that grasses primarily exploit shallow soil water whereas forbs access deeper layers, an expectation based largely on general rooting depth patterns and selected field studies (Schenk & Jackson, 2002; Nippert & Knapp, 2007). Our results show that water uptake depth is not reliably predicted by growth form in diverse temperate grassland communities.

High overlap was also observed at the plot scale with comparable representation of grasses and forbs, indicating that strong local overlap can occur despite significant differentiation among individual species. Water uptake may therefore be shaped more strongly by species identity and local environmental conditions than by broad herbaceous functional categories, as realized uptake depths integrates root distribution, soil moisture conditions, hydraulic traits and neighbour interactions (Freschet et al., 2021; Kühnhammer et al., 2020).

### Rooting systems as poor proxies for realized water use

Similarly, rooting system types provided only weak explanatory power for variation in water uptake depth. Although taprooted species tended to access slightly deeper layers in ALB and SCH, overlap with fibrous rooted species remained high, and differentiation was minimal in HAI. This weak correspondence reinforces the distinction between potential rooting architecture and realized water acquisition. Maximum rooting depth and root system morphology describe the soil volume that can potentially be explored, but the presence of roots at a given depth does not necessarily imply proportional water uptake from that depth (Nippert & Holdo, 2015; Guderle et al., 2018; Kühnhammer et al., 2020).

Thus, species with contrasting root architectures may still access similar water sources, while species with similar rooting systems may differ in water uptake depths. This helps explain the substantial overlap in water uptake observed among rooting system types in our study. However, this does not imply that trait-based prediction of water uptake depth should be abandoned, but rather suggests that more detailed root traits, particularly root distribution and hydraulic properties, may better predict water uptake depth than broad rooting system classifications.

### Implications for belowground coexistence mechanisms in temperate grasslands

Taken together, our findings suggest that belowground water use in temperate grasslands is characterized by fine scale species differentiation within a shared shallow soil domain rather than fixed vertical stratification among broad herbaceous functional groups (Fig. 4).

**Figure 4.**
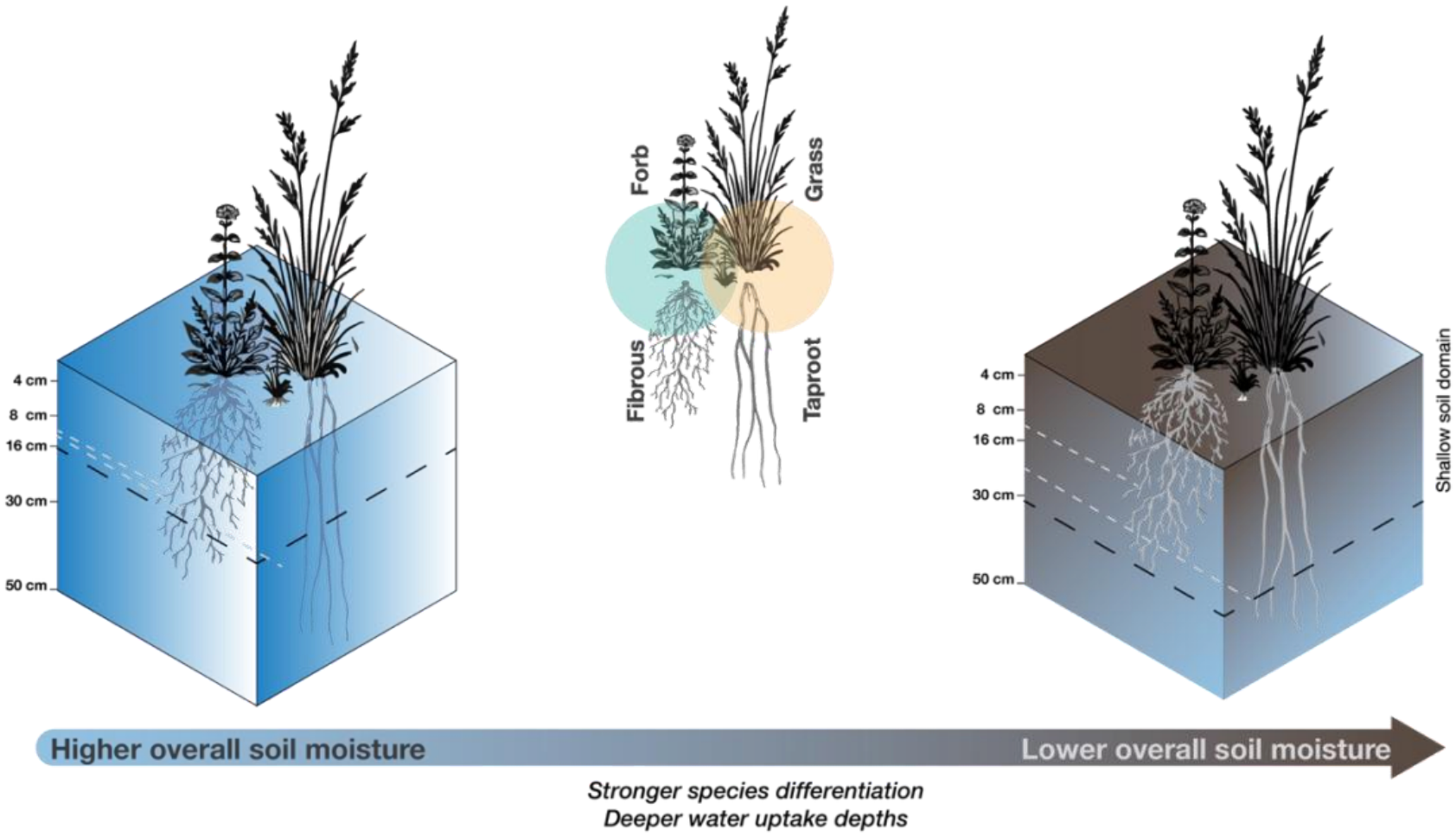
Conceptual synthesis of species differentiation in water uptake depth. Coexisting grassland species differentiate their water uptake depths within a shallow vertical zone in the soil profile, with greater differentiation and deeper uptake under lower overall soil moisture. Broad herbaceous functional classifications, show substantial overlap in water uptake depth.

By differing in where they acquire water, coexisting species may reduce overlap in resource use and potentially reduce interspecific competition, even when these differences occur within a relatively narrow vertical zone (Chesson, 2000; HilleRisLambers *et al*., 2012). The stronger differentiation observed under lower soil moisture further suggests that hydrological niche differentiation may become more pronounced as water becomes limiting. However, our study does not directly test whether these differences reduce competition or contribute to stable coexistence. Addressing this would require linking water uptake differentiation to species interactions, performance and demographic responses.

## Conclusion

This study demonstrates that coexisting grassland species consistently differ in water uptake depth, providing evidence for hydrological niche differentiation within a shared shallow soil zone. This differentiation becomes stronger under lower overall soil moisture, while broad herbaceous functional classifications based on growth form and rooting system capture little of the observed species level variation. These findings show that water uptake differentiation in temperate grasslands occurs at a fine spatial scale and is better captured at the species level than by broad herbaceous functional classifications.

## Acknowledgements

We thank the Plant Ecology group of the University of Tübingen for their valuable input and discussions throughout this project. We also thank the managers of the three Biodiversity Exploratories, Swabian Alb (Julia Bass, Max Müller), Hainich-Dün (Anna K. Franke), and Schorfheide-Chorin (Franca Marian, Max Müller, Uta Schumacher), and all former managers for their work in maintaining the plot and project infrastructure. We thank Victoria Grießmeier for giving support through the central office, and the respective data manager for managing the central database. We also thank Markus Fischer, Eduard Linsenmair, Dominik Hessenmöller, Daniel Prati, Ingo Schöning, François Buscot, Ernst-Detlef Schulze, Wolfgang W. Weisser, and the late Elisabeth Kalko for their role in setting up the Biodiversity Exploratories project. We thank the administration of the Hainich National Park, the UNESCO Biosphere Reserve Swabian Alb, and the UNESCO Biosphere Reserve Schorfheide-Chorin, as well as all landowners, for their excellent collaboration. The work has been funded by the German Research Foundation (DFG) as part of the Priority Programme 1374 “Biodiversity Exploratories” (DFG Ref No. MA9122/2-1). Fieldwork permits were issued by the responsible state environmental offices of Baden-Württemberg, Thuringia, and Brandenburg.

## Data availability

This experiment is based on data collected within the ExploreNiche project of the Biodiversity Exploratories program (DFG Priority Program 1374). All datasets used in this study are publicly available through the Biodiversity Exploratories Information System (BExIS) [https://doi.org/10.17616/R32P9Q]. The datasets are identified as: 32187 (Oxygen isotope content in plants from grassland plots in Germany, 2023) and 32186 (Oxygen isotope content in soil water from grassland plots in Germany, 2023). The authors strongly support the appropriate reuse and integration of these data in future research and database applications.

## Authors contributions

Maria Májeková conceived the idea. Maria Májeková and Sepideh Golshani designed the study. Sepideh Golshani, Tomáš Hájek, Undine Schöllkopf, Johanna Harrison, Barbora Rybová collected the data; Sepideh Golshani, Tomáš Hájek, Hassan Jafari, Maria Májeková, analyzed the data; Sepideh Golshani wrote the first draft; Katja Tielbörger, Pierre Liancourt, Maria Májeková contributed to writing.

## Supporting Information

**Figure S1.** Mixed effects models used to estimate plant water uptake depth from soil and plant δ¹⁸O values in the three study regions.

**Figure S2.** Map of the water uptake patterns across the three Biodiversity Exploratories regions in Germany.

**Figure S3.** Schematic of the isotope based estimation of water uptake depth and rain exclusion experiment.

**Figure S4.** Species specific regional differences in water uptake depth.

**Table S1.** Environmental, geological and geomorphological characteristics of the three Biodiversity Exploratories regions.

**Table S2.** List of sampled plant species and their occurrence across the three study regions.

**Table S3.** Species variation in water uptake depth across regions, based on estimated marginal means and pairwise regional contrasts.

**Table S4.** Effects of soil moisture and land use intensity on water uptake depth and mean pairwise differences among coexisting species.

