## Supporting information for "Species differentiate water uptake depths within a narrow vertical zone in temperate grasslands"

### Belowground niche partitioning in grasslands occurs within a narrow vertical zone

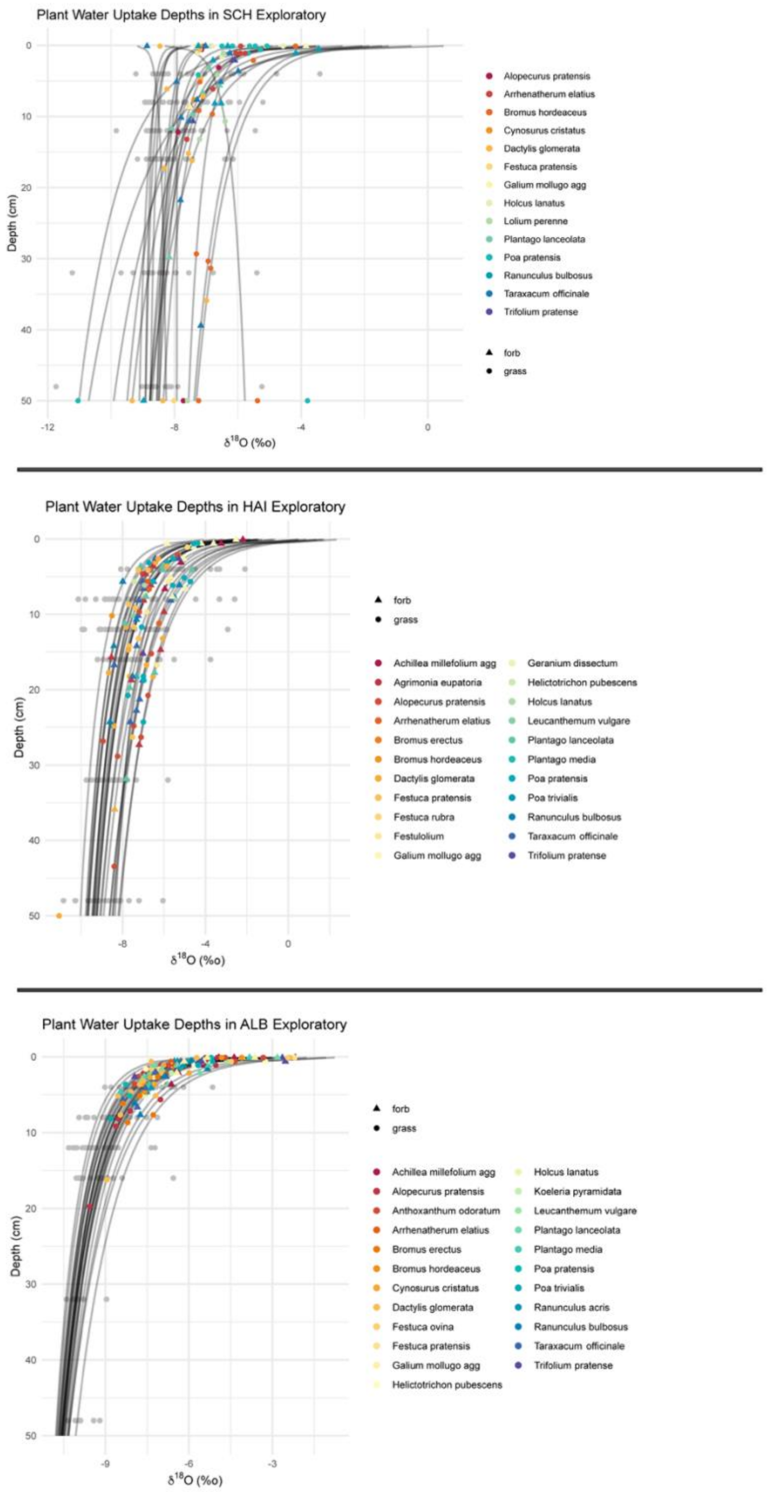


**Figure S1.** Mixed-effects models used to estimate plant water-uptake depth from $\delta^{18}O$ values in the three study regions.

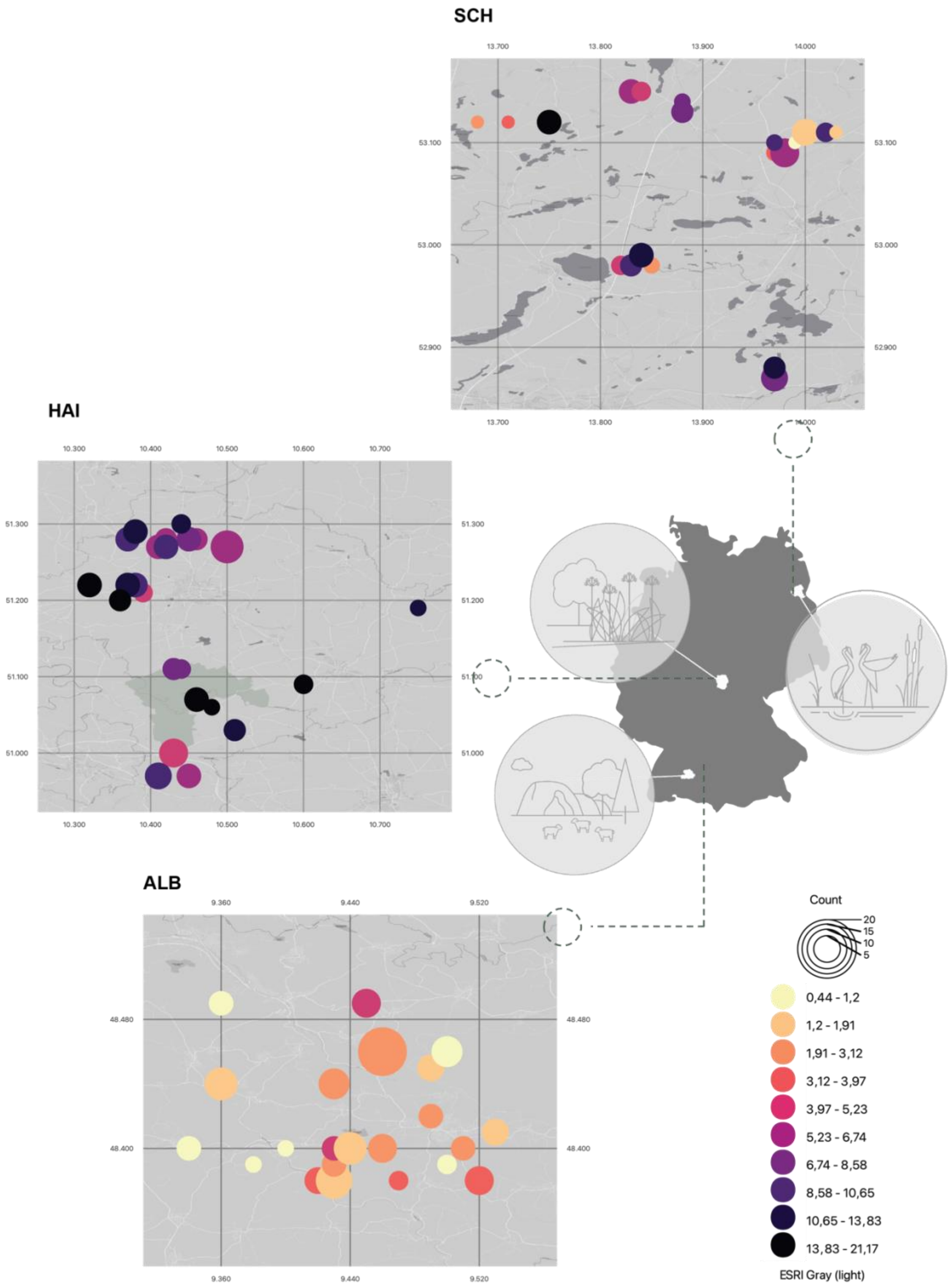


**Figure S2 Water uptake patterns across the three Biodiversity Exploratories regions in Germany.** Maps show Schwäbische Alb (ALB), Hainich Dün (HAI) and Schorfheide Chorin (SCH). Point colour indicates mean realized water uptake depth within each plot, while point size indicates the number of sampled species.

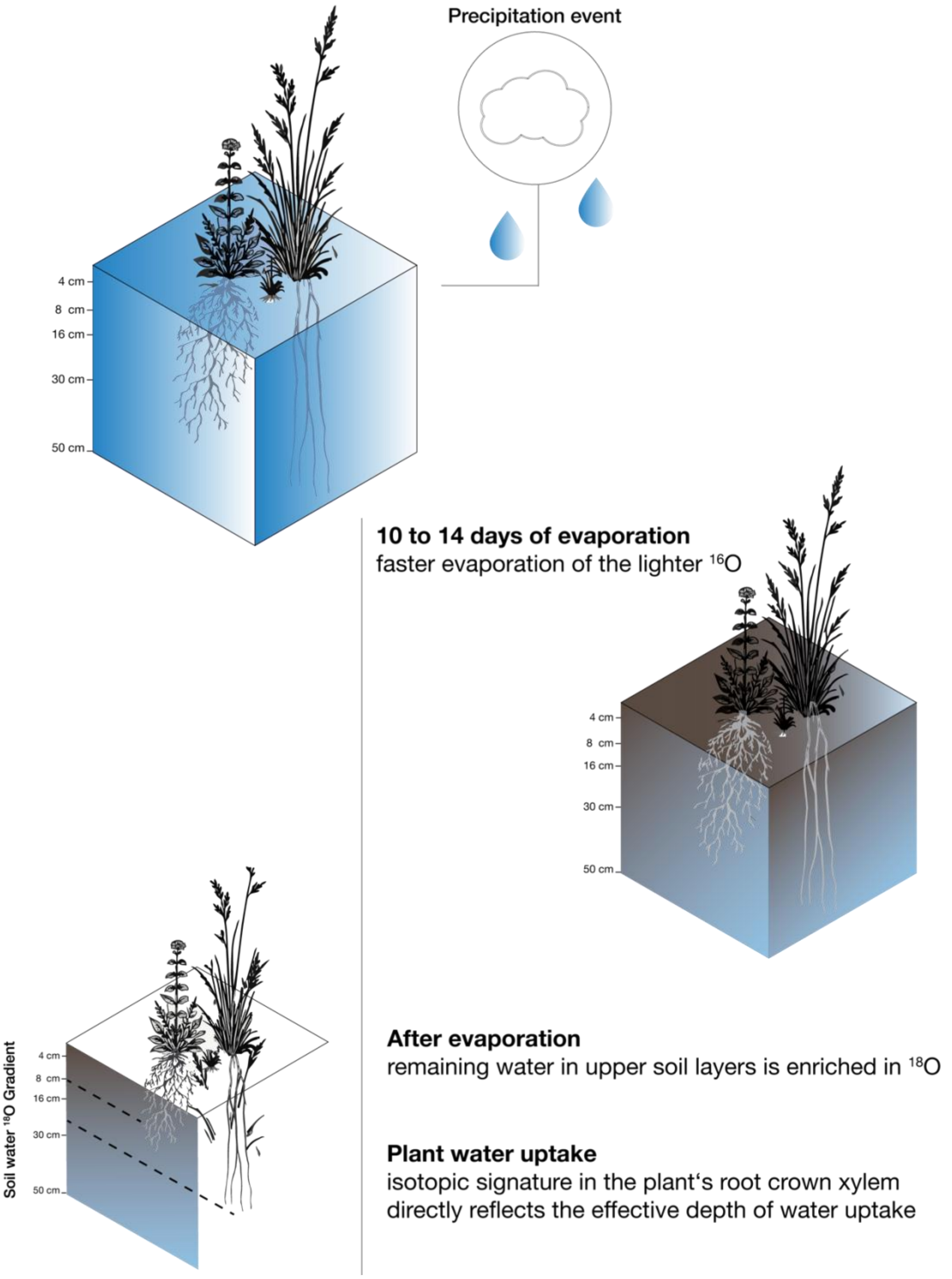


**Figure S3 Schematic of the rain exclusion and isotope based water uptake estimation.** Rainfall was excluded for 10–14 days to enhance vertical $\delta^{18}O$ gradients in soil water. Soil was sampled at multiple depths, and plant root crown $\delta^{18}O$ was compared with the soil isotope profile to estimate realized water uptake depth.

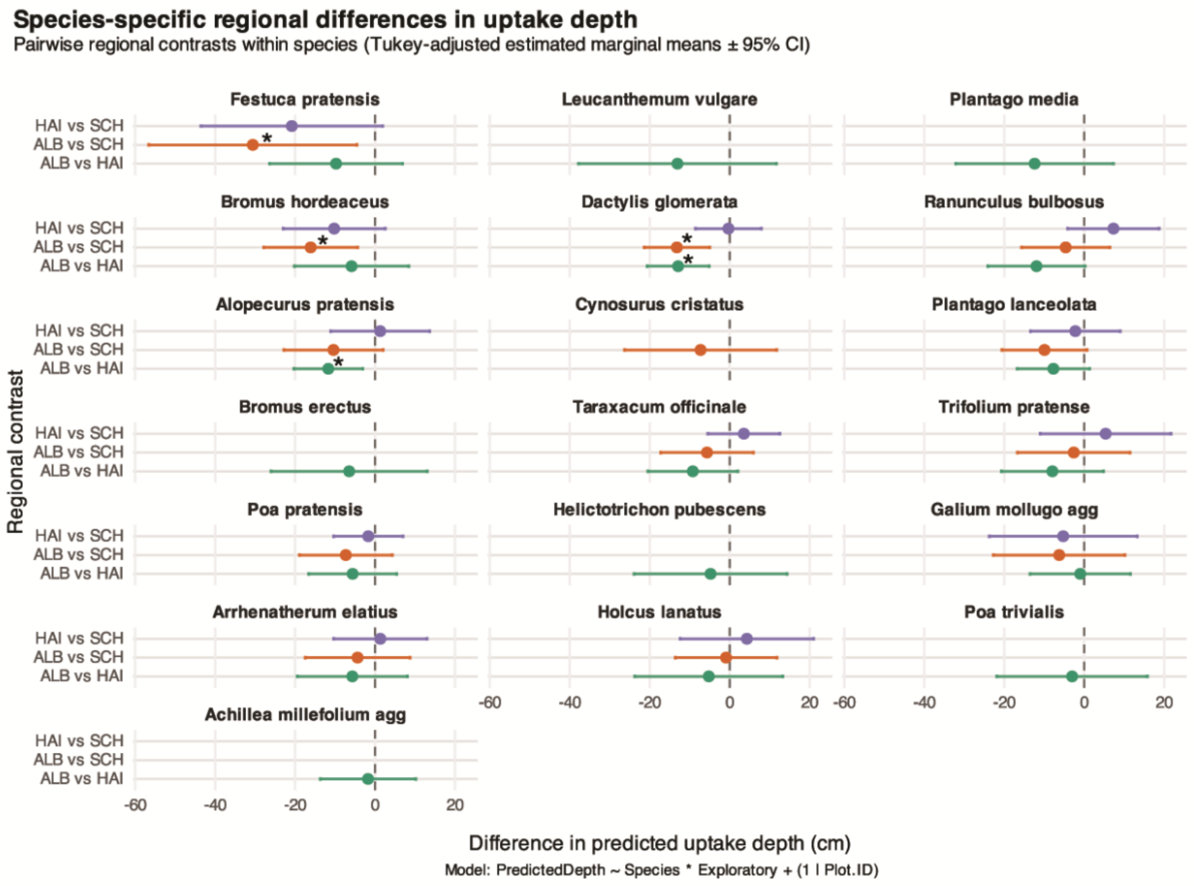


**Figure S4. Species-specific regional differences in water-uptake depth.** Pairwise contrasts of estimated marginal means (±95% CI) are shown for each species based on a linear mixed-effects model. Points represent Tukey-adjusted differences between regions; error bars indicate 95% confidence intervals. Intervals overlapping zero indicate no significant difference; significance levels are denoted by symbols (*)

**Table S1 Description of the three study regions within the German Biodiversity Exploratories.** Summary of environmental and geomorphological characteristics of the three. Variables include annual precipitation, elevation range, total area, dominant soil types, geological bedrock, and geomorphology (Fischer et al. 2010).

| *Variable* | *Schwäbische Alb (ALB)* | *Hainich-Dün (HAI)* | *Schorfheide-Chorin (SCH)* |
|---|---|---|---|
| ***Annual precipitation (mm)*** | 700–1000 | 500–800 | 480 (East) – 580 (West) |
| ***Elevation (m a.s.l.)*** | 460–860 | 285–550 | 3–140 |
| ***Area (km²)*** | ~422 | ~130 | ~1300 |
| ***Dominant soil types*** | Leptosols (steep slopes), Cambisols | Eutric Cambisols, Stagnosols (on loess over Triassic limestone), Vertisols | Histosols (drained), Luvisols, Gleysols, Cambisols, Albeluvisols |
| ***Geological bedrock*** | Loess over calcareous Jurassic shell limestone; grasslands rich in clay | Calcareous bedrock with karst features | Glacial till, often covered by glaciofluvial or aeolian sands; fertile loamy soils in grasslands |
| ***Geomorphology*** | Mid-mountain range | Hilly terrain (Central German hilly lands) | Plain to gently undulating moraine landscape with fens and wetlands in depressions |

**Table S2 Selected grassland species.** List of the plant species sampled in each region. Species were chosen based on their high local abundance and occurrence across multiple plots and regions.

| Species | ALB | HAI | SCH |
|---|---|---|---|
| Achillea millefolium agg. | x | x | |
| Agrimonia eupatoria | | x | |
| Alopecurus pratensis | x | x | x |
| Anthoxanthum odoratum | x | | |
| Arrhenatherum elatius | x | x | x |
| Bromus erectus | x | x | |
| Bromus hordeaceus | x | x | x |
| Cynosurus cristatus | x | | x |
| Dactylis glomerata | x | x | x |
| Festuca ovina | x | | |
| Festuca pratensis | x | x | x |
| Festuca rubra | | x | |
| Festulolium | | x | |
| Galium mollugo agg. | x | x | x |
| Geranium dissectum | | x | |
| Helictotrichon pubescens | x | x | |
| Holcus lanatus | x | x | x |
| Koeleria pyramidata | x | | |
| Leucanthemum vulgare | x | x | |
| Lolium perenne | | | x |
| Plantago lanceolata | x | x | x |
| Plantago media | x | x | |
| Poa pratensis | x | x | x |
| Poa trivialis | x | x | |
| Ranunculus acris | x | | |
| Ranunculus bulbosus | x | x | x |
| Taraxacum officinale | x | x | x |
| Trifolium pratense | x | x | x |

**Table S3 Species variation in water-uptake depth across regions.** Estimated marginal means (emmean; cm) of plant water-uptake depth for each species across regions, derived from a linear mixed-effects model (PredictedDepth ~ Species × Exploratory + (1 | Plot.ID)). Species present in ≥2 regions were included. The minimum adjusted p-value (min p) denotes the strongest pairwise regional contrast per species; regional shift indicates at least one significant contrast ($p < 0.05$). Depth ranking (shallow → deep) is based on estimated marginal means within species.

| *Species* | *ALB (cm)* | *HAI (cm)* | *SCH (cm)* | *Depth ranking (shallow → deep)* | *Regional shift* | *min p* |
|---|---|---|---|---|---|---|
| ***Dactylis glomerata*** | 3.51 | 16.35 | 16.79 | ALB < HAI ≈ SCH | Yes | **0.0003** |
| ***Alopecurus pratensis*** | 4.37 | 15.96 | 14.63 | ALB < HAI ≈ SCH | Yes | **0.0045** |
| ***Bromus hordeaceus*** | 1.91 | 7.76 | 17.87 | ALB < HAI < SCH | Yes | **0.0040** |
| ***Festuca pratensis*** | 1.04 | 11.18 | 30.47 | ALB < HAI < SCH | Yes | **0.0164** |
| *Arrhenatherum elatius* | 2.18 | 8.07 | 6.64 | ALB < SCH < HAI | No | >0.05 |
| *Galium mollugo agg* | 0.63 | 1.40 | 7.02 | ALB < HAI < SCH | No | >0.05 |
| *Holcus lanatus* | 1.78 | 8.00 | 2.88 | ALB < SCH < HAI | No | >0.05 |
| *Plantago lanceolata* | 2.16 | 10.03 | 12.59 | ALB < HAI < SCH | No | >0.05 |
| *Poa pratensis* | 2.08 | 7.40 | 9.12 | ALB < HAI < SCH | No | >0.05 |
| *Ranunculus bulbosus* | 3.37 | 15.08 | 7.62 | ALB < SCH < HAI | No | 0.055 |
| *Taraxacum officinale* | 3.58 | 13.08 | 9.21 | ALB < SCH < HAI | No | >0.05 |
| *Trifolium pratense* | 0.94 | 8.94 | 2.70 | ALB < SCH < HAI | No | >0.05 |
| *Achillea millefolium agg* | 0.31 | 2.18 | — | ALB < HAI | No | >0.05 |
| *Bromus erectus* | 5.12 | 11.20 | — | ALB < HAI | No | >0.05 |
| *Cynosurus cristatus* | 1.60 | — | 9.03 | ALB < SCH | No | >0.05 |
| *Helictotrichon pubescens* | 1.63 | 5.59 | — | ALB < HAI | No | >0.05 |
| *Leucanthemum vulgare* | 1.35 | 14.65 | — | ALB < HAI | No | >0.05 |
| *Plantago media* | 2.19 | 14.23 | — | ALB < HAI | No | >0.05 |
| *Poa trivialis* | 1.41 | 4.27 | — | ALB < HAI | No | >0.05 |

**Table S4 Effects of soil moisture and land use intensity on realized water uptake depth and species differentiation.** For realized water uptake depth, a linear mixed effects model was fitted to log transformed predicted uptake depth, with species and plot as random intercepts. Species differentiation was quantified as the mean pairwise difference in log water uptake depth within plots and analysed using a linear model. Both models included overall soil moisture, vertical soil moisture distribution, land use intensity (LUI), and region simultaneously.

| *Response* | *Predictor* | *Estimate* | *SE* | *df* | *t* | *p* |
|---|---|---|---|---|---|---|
| **Log water uptake depth** | Overall soil moisture | −0.035 | 0.006 | 79.39 | −6.26 | **<0.001** |
| | Vertical soil moisture distribution | 0.175 | 0.179 | 60.76 | 0.98 | 0.332 |
| | **LUI** | 0.309 | 0.165 | 60.15 | 1.87 | 0.067 |
| | HAI vs ALB | 1.425 | 0.275 | 63.33 | 5.18 | **<0.001** |
| | SCH vs ALB | 0.682 | 0.287 | 68.70 | 2.38 | **0.020** |
| **Mean pairwise difference in log uptake depth** | Overall soil moisture | −0.010 | 0.004 | 66 | −2.26 | **0.027** |
| | Vertical soil moisture distribution | −0.056 | 0.153 | 66 | −0.37 | 0.715 |
| | **LUI** | 0.129 | 0.142 | 66 | 0.91 | 0.368 |
| | HAI vs ALB | −0.390 | 0.234 | 66 | −1.67 | 0.100 |
| | SCH vs ALB | −0.038 | 0.237 | 66 | −0.16 | 0.874 |

**Land use intensity calculation**

Land use intensity (LUI) data for 2023 were obtained from the Biodiversity Exploratories land use dataset. LUI integrates fertilization, mowing and grazing intensity following Blüthgen et al. (2012). Fertilization intensity was quantified as the amount of nitrogen applied (kg N $ha^{-1}$ $yr^{-1}$), mowing intensity as the number of cuts per year, and grazing intensity as livestock unit grazing days $ha^{-1}$ $yr^{-1}$. Each component was standardized relative to its mean within the corresponding Exploratory region, and the standardized components were combined into the compound LUI index.
